# Skeletal muscle-derived Human Mesenchymal Stem Cells: influence of different culture conditions on proliferative and myogenic capabilities

**DOI:** 10.1101/2020.05.12.090746

**Authors:** Stefano Testa, Carles Sanchez Riera, Ersilia Fornetti, Federica Riccio, Claudia Fuoco, Sergio Bernardini, Jacopo Baldi, Marco Costantini, Maria Laura Foddai, Stefano Cannata, Cesare Gargioli

## Abstract

Skeletal muscle tissue is characterized by restrained self-regenerative capabilities, being ineffective in relation to trauma extension both in time span (e.g. chronic diseases) and in size (e.g. large trauma). For these reasons, tissue engineering and/or cellular therapies represent a valuable solution in the cases where the physiological healing process failed. Satellite cells, the putative skeletal muscle stem cells, have been the first solution explored to remedy the insufficient self-regeneration capacity. Nevertheless, some limitation related to donor age, muscle condition, expansion hitch and myogenic potentiality maintenance have limited their use as therapeutic tool. To overcome this hindrance, different stem cells population with myogenic capabilities have been investigated to evaluate their real potentiality for therapeutic approaches, but, as of today, the perfect cell candidate has not been identified yet.

In this work, we analyze the characteristics of skeletal muscle-derived human Mesenchymal Stem Cells (hMSCs), showing the maintenance/increment of myogenic activity upon differential culture conditions. In particular, we investigate the influence of a commercial enriched growth medium (Cyto-Grow), and of a medium enriched with either human-derived serum (H.S.) or Platelet-rich Plasma (PrP), in order to set up a culture protocol useful for employing this cell population in clinical therapeutic strategies. The presented results reveal the remarkable effects of H.S. in the enhancement of hMSC proliferation and myogenic differentiation.

## 1 Introduction

The self-regenerative process of skeletal muscle tissue is a complex phenomenon that engages several types of resident and circulating stem cells with different potentialities (Yin et al., 2013; Čamernik et al., 2018). Among all of these cell types, the most important involved in the repairing process are satellite (Mauro, 1961) and non-satellite cell populations (Crisan et al., 2008; Sacchetti et al., 2016). While the former has been well characterized in the last decades and their activation mechanism unraveled relying on satellite cell niche regulation by the complex interaction among Notch, Collagen V and Calcitonin receptor (Baghdadi et al., 2018), the latter are gaining increasing interest in the research community thanks to readiness of isolation and expansion, and last but not least the better migratory capacity (Rando et al., 1995; Ferrari et al., 1998; Sacco et al., 2008).

Notably, non-satellite cell population is a heterogeneous group of mesenchymal stem cells (MSCs) that includes interstitial cells called PW1^+^/Pax7^-^ Interstitial Cells (PICs) (Mitchell et al., 2010), fibro-adipogenic progenitors (FAPs) (Uezumi et al., 2010), muscle side population cells (SP) and muscle resident pericytes (Gussoni et al., 1999; Kumar et al., 2017).

MSCs were initially described in the bone marrow as bone marrow-derived mesenchymal stromal/stem cells (BM-MSCs) for their unique combination of features, which include fibroblast-like morphology, clonogenicity, multipotency, and *in vitro* adherence on plastic surface unlike the hematopoietic counterpart (Friedenstein et al., 1970; Pittenger et al., 1999). Cells with similar *in vitro* abilities have been isolated from numerous adult tissues and organs, including skeletal muscle, both from small mammals and human biopsies (Hass et al., 2011; Čamernik et al., 2018). So far, the most popular and studied tissue source employed for MSC isolation has been the bone marrow thanks to its availability and accessibility in the human body. Nevertheless, over the last decade other tissue sources have been explored such as fat tissue, umbilical cord, dental pulp, skin, placenta and even brain (Yamada et al., 2010; Appaix, 2014; Orciani et al., 2014; Li et al., 2015; Bieback and Netsch, 2016; Pelekanos et al., 2016; Amati et al., 2017). Despite heterogeneity primarily due to different isolation tissue sources, MSCs maintain characteristic expression markers such as CD90, CD44, CD73, CD29, and CD105 while missing the hematopoietic ones such as CD34, CD45 and CD11 (Haynesworth et al., 1992; Lodie et al., 2002; Suva et al., 2004). Besides, they retain and share similar differentiation potential in mesoderm-lineage tissues including bone, fat, cartilage and skeletal muscle (Okamoto et al., 2002; Sottile et al., 2002; Zhang et al., 2009; Almalki and Agrawal, 2016; Kozlowska et al., 2019).

As regards skeletal muscle tissue, bone marrow-derived MSCs differentiate into myogenic lineage exclusively upon exposure to demethylating agent 5-azacytidine (Wakitani et al., 1995; Jackson et al., 2007) or in co-culture with myocytes (Lee et al., 2005). More recently, Sacchetti and collaborators have demonstrated that skeletal muscle-derived MSCs have, beyond an intrinsic heterogeneity, spontaneous and important myogenic capabilities *in vitro* and *in vivo*. Moreover, the Authors have shown that the differentiation potentiality may vary radically according with the tissue origin and that MSCs immune-profile does not reflect identical cells and function (Sacchetti et al., 2016). Anyway, human MSCs (hMSCs), due to their multiple potentialities, could still represent a good candidate for cell therapy (De Bari et al., 2003; Oppermann et al., 2014; Klimczak et al., 2018; Pittenger et al., 2019). Hence, in order to translate MSCs into the actual clinical scenario, researchers still need to address a long-standing challenge: to produce *in vitro* a clinically relevant number of cells without affecting their differentiation capacity. Moreover, culture conditions should be standardized to fulfill good manufacturing practice (GMP) protocols and to avoid possible contamination or immunological reactions due to xenogeneic medium supplement, e.g. animal derived serums (Tonti and Mannello, 2008). In fact, as demonstrated in numerous studies, the expansion of hMSCs strongly depends on the culture conditions, being anchorage-dependent and requiring medium supplemented with 10-20% serum (De Bari et al., 2003; Meuleman et al., 2006; Čamernik et al., 2018; Musiał-Wysocka et al., 2019). Additionally, interactions among cells, growth surface and surrounding medium influence many aspects of cell behavior, such as efficiency of isolation, proliferation rate, maintenance in culture, stemness and differentiation potentiality (Kozlowska et al., 2019; Musiał-Wysocka et al., 2019).

Given all these key aspects, over the past years a growing interest has been focused on biologic agents such as blood derivatives such as Platelet-rich plasma (PrP) to complement the cell culture medium and/or significantly ameliorate musculoskeletal tissue healing (Hamid et al., 2014; Kunze et al., 2019). However, their real benefic effect is still questioned (Grassi et al., 2018).

In order to shed some lights on the potentialities of skeletal muscle-derived human mesenchymal stem cells for skeletal muscle regeneration, here we compare different culture conditions evaluating immunophenotypical aspect, cell growth and myogenic differentiation capability

## Materials and Methods

### 2.1 Isolation of mesenchymal stem cells from muscle biopsies and cell culture

hMSCs were isolated from skeletal muscle tissue using a protocol that includes mechanical mincing, enzymatic digestion with type II collagenase, filtration, and selection of the colonies on plastic surface at low confluence (Sacchetti et al., 2016; Vono et al., 2016). Briefly, human skeletal muscle biopsies are finely minced with a surgical knife and collected in a solution of collagenase type II (100 U/ml in PBS Ca2+/ Mg2+), subsequently left to incubate in a thermal shaker for 45 minutes at 37 °C. After digestion, the solution is centrifuged at 300g for 10 minutes at room temperature and then aspirated without disturbing the pellet. The pellet is then resuspended in 15 ml of PBS and filtered through progressively finer cell strainers: 100 µm, 70 µm and 40 µm. Cells are counted with a Burker counting chamber and plated on conventional Petri dishes (BD Falcon) at low confluence (10^3^ cells/cm^2^) to promote the growth of cells that have clonogenicity and therefore stem potentiality. The freshly isolated mesenchymal stem cells were divided into two experimental groups respectively cultured in either Alpha-MEM (Gibco) supplemented with 20% heat-inactivated fetal bovine serum (FBS, EuroClone), penicillin (100 IU/mL, Gibco) and streptomycin (100 mg/mL, Gibco), or Cyto-grow medium supplemented with penicillin (100 IU/mL, Gibco) and streptomycin (100 mg/mL, Gibco). In both conditions, cells are cultured at 37°C and 5% CO^2^ and left to incubate for 15 days, time required for colonies formation. Alternatively, isolated cells are harvested and used for flow cytometry analysis.

### 2.2 Flow cytometry analysis

1×10^6^ cells for each experimental condition are harvested with Lonza™ Trypsin-Versene™-Trypsin-EDTA (Fisher Scientific, #BE17-161E), resuspended and centrifuged at 300g for 10 minutes at room temperature. Cells are washed twice with PBS supplemented with BSA 0,5% (Bovine Serum Albumin, AppliChem, #A1391) and EDTA 2mM at 300g for 10 minutes at 4*°*C. Samples are then resuspended in PBS and incubated with APC-A700 anti-human CD56 (N901, #B92446 Beckman Coulter), APC-A750 anti-human CD90 (Thy-1/310, ***#***B36121 Beckman Coulter) and Vioblue anti-human CD45 antibodies (REA747, #5180719178 Miltenyi Biotec) for 30 minutes at 4°C. After incubation, cells are washed in PBS, centrifuged at 300g for 10 minutes at 4*°*C and resuspended in PBS. Samples are visualized on Cytoflex S, 3 lasers (488 nm, 405 nm and 638 nm) and 13 detectors (Beckman Coulter). Live cells were gated based on side scatter and forward scatter. Data are analyzed by CytExpert software (Beckman Coulter).

### 2.3 Growth curves

Growth curves were obtained using 48-well plates containing 2×10^4^ cells/well. Well plates were prepared for every experimental point (2, 5, 7 and 9 days) and every experimental condition: i) Alpha-MEM supplemented with 20% FBS (control), ii) 5% human serum (low), iii) 10% human serum (medium) or iv) 20% human serum (high). Same procedure was followed to test the effect of the serum donor age on cell proliferation. In this case experimental conditions were: Alpha-MEM supplemented with 20% of human serum from donors in the age range of: i) 20-30 (young), ii) 31-50 (adult) or iii) 51-65 (senior) years old. In the case of PrP assay the experimental conditions were: Alpha-MEM supplemented with 20% FBS (control) and with 5×10^5^ platelets/ml (low), 1×10^6^ platelets/ml (medium) or 1,5×10^6^ platelets/ml (high). Cellular proliferation was evaluated by harvesting cells at each time point and scoring the media in a Burker counting chamber.

### 2.4 Immunofluorescence analysis

Cells were fixed with 4% PFA in PBS for 10 minutes at 4°C and processed for immunofluorescence analysis as previously described (Testa et al., 2017). Briefly, cells are washed with PBS and blocked with 10% goat serum in PBS for 1 h at room temperature (RT). Subsequently, cells are incubated with the primary antibody anti-myosin heavy chain (MF20, mouse monoclonal, DHSB, diluted 1:2) or anti-ki67 (rabbit polyclonal, Novus Biologicals #NB110-89717, diluted 1:200) for 1 h, followed by incubation with respectively Alexa Fluor 555-conjugated goat anti-mouse IgG (H+L) (Thermo Fisher Scientific #A21422, diluted 1:400) and 488-conjugated goat anti-rabbit IgG (H+L) (Thermo Fisher Scientific #A11008, diluted 1:400) for 1 h. Finally, nuclei are stained with 300 nM DAPI (Thermo Fisher Scientific) for 10 min. Specimens were viewed using a Nikon TE 2000 epifluorescence microscope equipped with a Photometrics Cool SNAP MYO CCD camera.

### 2.5 Statistical analysis

All experiments were performed in biological and technical triplicate (*n*=9). Data were analyzed using GraphPad Prism 7, and values are expressed as means ± standard error (SEM). Statistical significance was tested using ONE WAY ANOVA test. A probability of less than 5% (p<0,05) was considered to be statistically significant.

## Results

### 3.1 Influence of different culture conditions on stemness and myogenic capability of skeletal muscle-derived hMSCs

Mononucleated cells from human derived skeletal muscle biopsies were examined at different time points by flow cytometry analysis in order to test whether different culture media would affect cell populations heterogeneity and behavior. In this regard, hMSCs were divided in two experimental groups according to the differential culture conditions: i) Alpha-MEM supplemented with 20% of FBS (standard medium) or ii) Cyto-grow (rich medium). The two experimental groups were analyzed at different time points (t0-t3) starting from isolation up to late doubling time, namely: total mononucleated cells soon after isolation from fresh muscle biopsy (**t0**), colonies formation stage according to MSCs conduct (after 15 days on standard plastic culture) (Sacchetti et al., 2007, 2016) (**t1**), expansion passage 3 (**t2**) and expansion passage 9 (**t3**) (**Figure 1A**). Cells were, accordingly, analyzed by flow cytometry analysis to evaluate the expression of hMSC stemness marker CD90 and the human myogenic marker CD56, further using CD45 as a negative marker (hematopoietic compartment). Analysis at t0 revealed a heterogeneous population of mononucleated cells, presenting several stem cells CD90^+^ (15,75%) together with abundant myogenic cells CD56^+^ (76,65%) displayed in the dot-plot, besides hematopoietic stem cells positive for CD45 (11,45%) (**Figure 1A, t0**). Hence, t0 population has been split and cultured in two different media (Alpha-MEM or Cyto-grow, used in the following stages) revealing differences in terms of marker expression already at **t1**. Here, the colonies formed in standard medium presented the tendency to segregate in two subpopulations: one exclusively CD90^+^ and the other one double positive CD56^+^/CD90^+^. Differently, cells cultured in rich medium formed a more homogeneous double positive population CD56^+^/CD90^+^ (**Figure 1A t1**). At **t2**, in standard medium, two well distinct cell subpopulations became more evident reaching at **t3** 29,22% of CD90^+^ cells and 70,65% of double positive CD56^+^/CD90^+^ cells, while expansion in rich medium selected myogenic CD56^+^ cell population at the expenses of CD90^+^/CD56^+^ double positive one (**Figure 1A, t2 and t3**). The CD45^+^ hematopoietic stem cells are lost on both groups already at **t1 (Figure 1A)**. Immunofluorescence against muscle terminal differentiation marker Myosin Heavy Chain (MyHC) has also been performed at **t3** on both experimental groups, revealing in both cases a remarkable spontaneous myogenic capabilities of skeletal muscle-derived hMSCs (**Figure 1B**).

**Figure 1.**
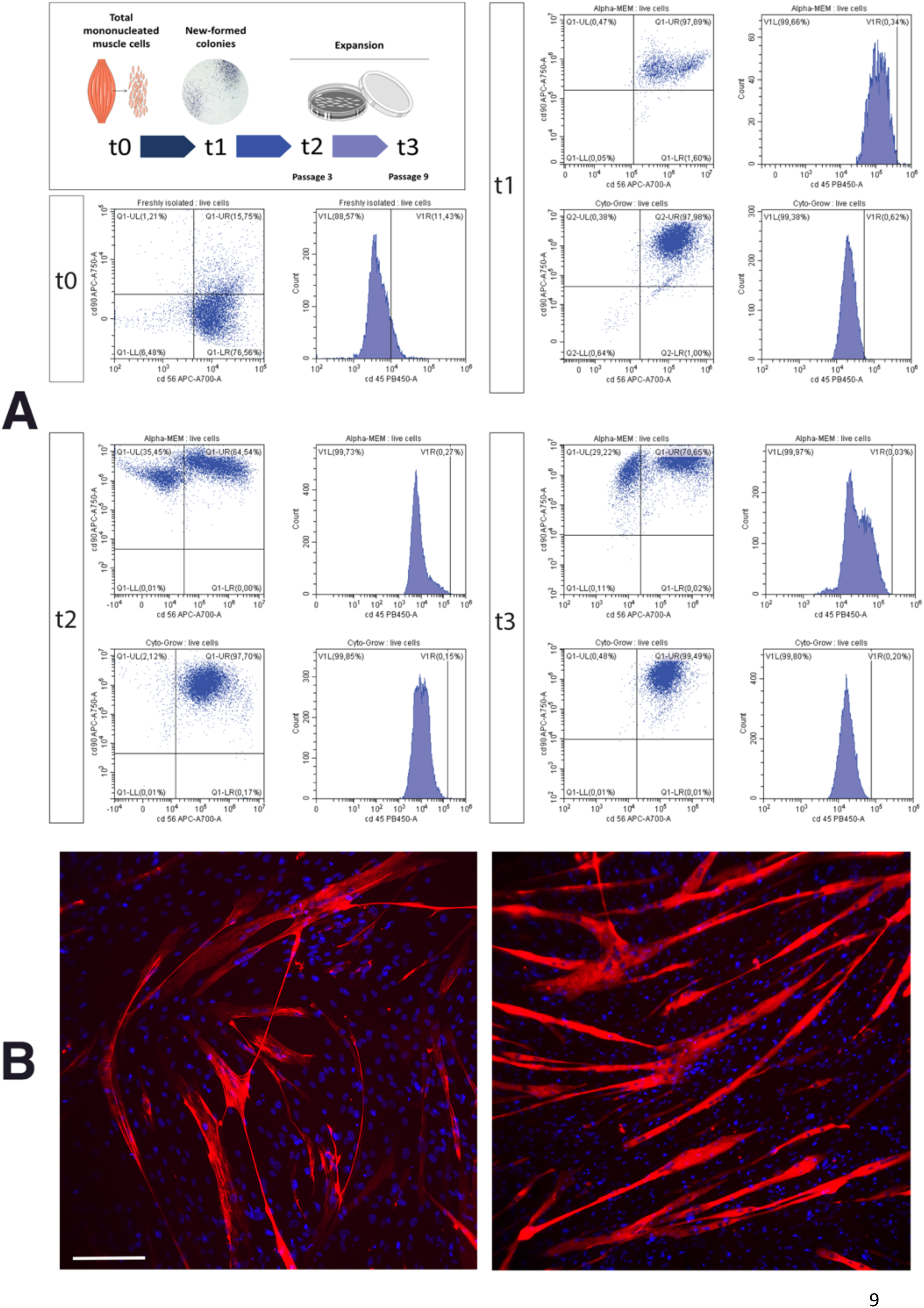
hMSC characterization by flow cytometric analysis. A) Scheme representing the time points analyzed in the flow cytometric analysis, from skeletal muscle-derived hMSCs isolation up to late doubling time: total mononucleated cells freshly isolated from the tissue (t0), new formed colonies after 15 days (t1), expansion passage 3 (t2) and expansion passage 9 (t3). Scatter dot plots showing hMSC population analyzed by flow cytometry for CD56 and CD90 in relation to different culture media: alpha-MEM (top) and Cyto-Grow (down). The histograms on the right of the dot plots (in each time points) represent CD45 analysis. B) Immunofluorescence staining with anti-myosin heavy chain (red) on terminally differentiated hMSCs (15 days) cultured in Alpha-MEM (left) and Cyto-Grow (right); nuclei were counterstained with DAPI (blue). Scale bar 150μm.

### 3.2 Employment of different human serum concentration as medium supplement for hMSC cell culturing

In order to test whether human serum could be employed as medium supplement for hMSC growth and thus satisfy the need of MSC culture standardization avoiding animal derivatives, different concentration of this compound has been used to investigate its efficacy. Therefore, hMSCs have been cultured in Alpha-MEM supplemented with 20% FBS as standard (control) medium, or at increasing concentration of human serum: low (5%), medium (10%) and high (20%). At different time points (2, 5, 7 and 9 days of culture) the cells were harvested to score proliferation rate. hMSCs growth revealed a significant dependence on human serum concentration: in particular low concentration (5%) of human serum were comparable to the control medium (20% FBS) growth rate, while already at medium concentration (10%) it was possible to observe a remarkable enhancement in the hMSC proliferation, reaching a statistical significant increment at high concentration (20%) in every analyzed time point up to a more than 5-fold increase at day 9 (**Figure 2A**).

**Figure 2.**
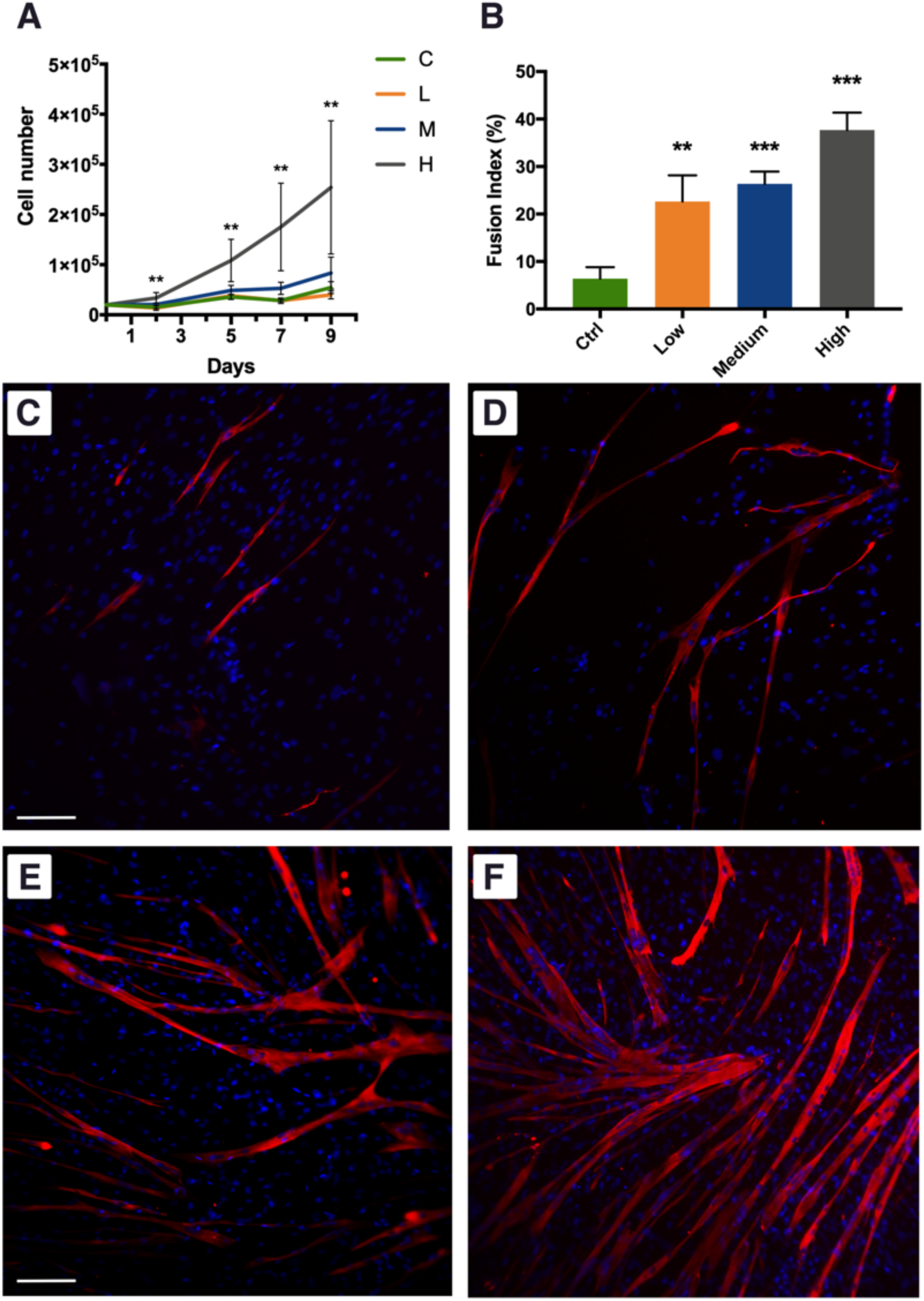
Effect of human serum on hMSC proliferation and differentiation. A) Histogram representing hMSC fusion index at day 9 of culturing in the investigated conditions. B) hMSC proliferation curve analyzed upon the different medium supplementation exposure. Statistics: One-way Anova test. p > 0.05^*^, p > 0.01^**^, p > 0.001^***^ (n= 3). C-F) Anti MyHC immunolabelling on hMSCs cultured with different supplemented Alpha-MEM media: (C) control (20% FBS), (D) Low (5% Human Serum), (E) Medium (10% Human Serum) and (F) High (20% Human Serum). Nuclei were stained by DAPI (Blue). Scale bars 100 μm.

Myogenic differentiation has been assessed by MyHC expression by means of immunofluorescence analysis at culturing day 9, a suitable time allowing cells to fuse and differentiate into myotubes. A clear trend was also observed for the number of formed myotubes, with the highest number of myotubes obtained with the highest human serum concentration (**Figure 2C-F**). These observations have been further confirmed by the fusion index analysis, a quantitative indicator of the cell differentiation level that is calculated by dividing the number of nuclei present in the myotubes (threshold: 3 nuclei per myotube) by the number of total nuclei. This analysis further confirmed that a better cell differentiation can be obtained using high level of human serum reaching the maximum value (4-fold increase compared to control) at 20% (**Figure 2B**).

### 3.3 Human serum age-dependence on hMSC cultures

Observing the remarkable influence of human serum concentration on hMSC proliferation and being the higher concentration (20%) the more effective in enhancing cell growth rate and differentiation, we additionally investigated whether the age of the donor-derived serum would affect this capability. Accordingly, hMSCs have been cultured for 9 days in standard medium supplemented with 20% of human serum from young (20-30 years old), adult (31-50 years old) or senior (51-65 years old) donors. At different experimental time points (2, 5, 7 and 9 days), cells exposed to different donor-age sera were harvested and counted to evaluate the proliferation rate. The resulting growth curves revealed no significantly differences among the three conditions (**Figure 3**), suggesting that the increase of hMSC proliferation rate is not dependent on the age of the donor-derived serum.

**Figure 3.**
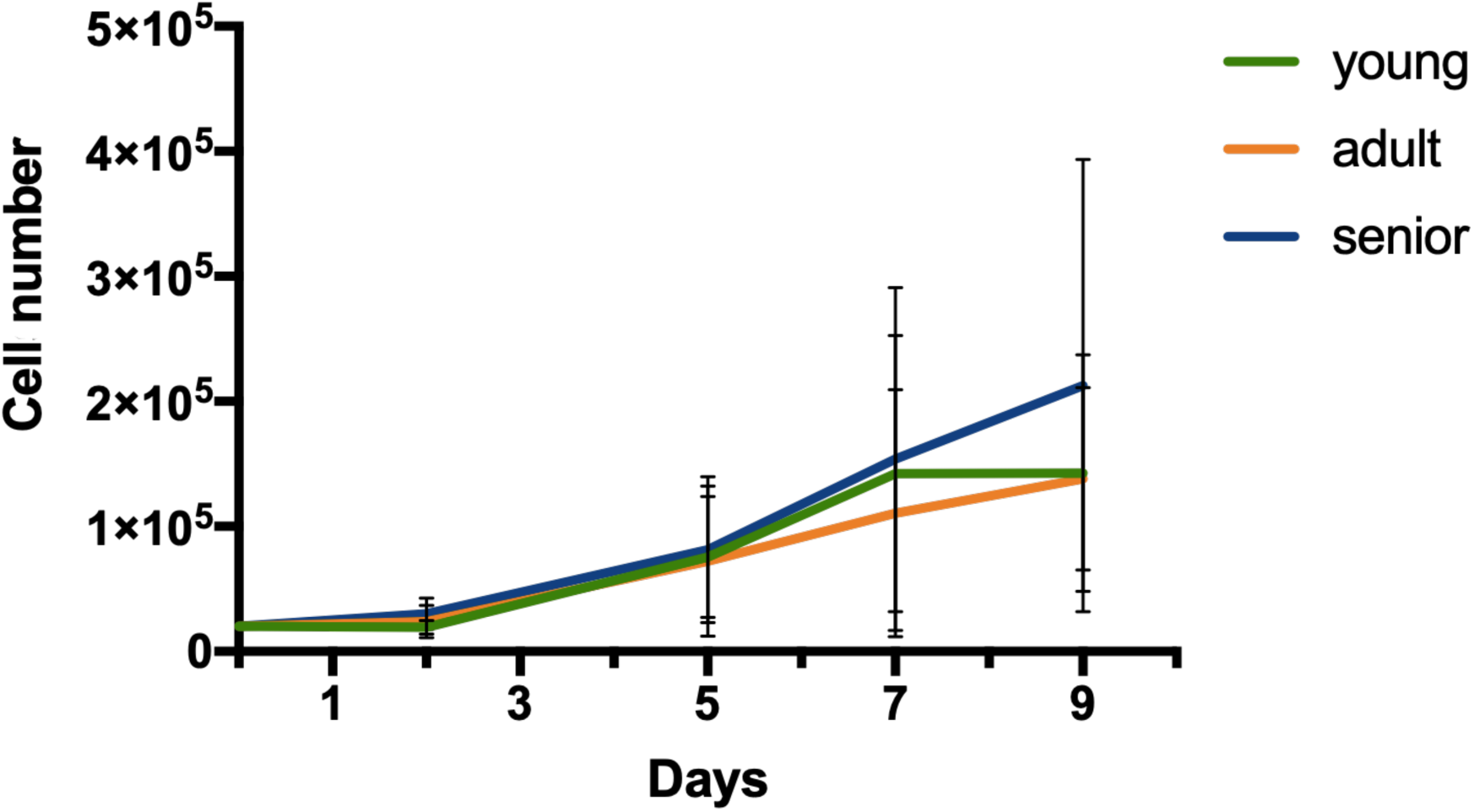
Different donor-age derived sera do not affect hMSC proliferation rate. Graph representing hMSC cell proliferation upon exposure to 20% human serum from different groups of donor-age: young (20-30 years old), adult (31-50 years old) and senior (51-65 years old). Statistics: One-way Anova test. p > 0.05^*^, p > 0.01^**^, p > 0.001^***^ (n= 3).

### 3.4 Influence of different Platelet-rich plasma (PrP) concentrations on hMSC proliferation

In order to test PrP effect on cell division rate, hMSCs have been cultured with low (5×10^5^ platelets/ml), medium (1×10^6^ platelets/ml) and high (1,5×10^6^ platelets/ml) concentrations of PrP as medium supplement. Cells were harvested and counted at 2, 5, 7 and 9 days upon PrP exposure. The resulting growth curves revealed that medium supplementation with high concentration of PrP significatively increased cell division for 7 days, while a decrease was observed at day 9 (**Figure 4A**). The other concentrations of PrP demonstrated a relative moderate and progressive increase up to day 7, reaching the plateau at day 9. The PrP effect on hMSC proliferation has been further confirmed by immunofluorescence analysis for Ki67 (proliferation marker) at day 9 (**Figure 4C-F**). The obtained results, plotted scoring Ki67^+^ nuclei, were consistent with the cell growth curves showing a significant increase at low and medium PrP concentrations, and a soft decrease at high concentration exposure (**Figure 4B**).

**Figure 4.**
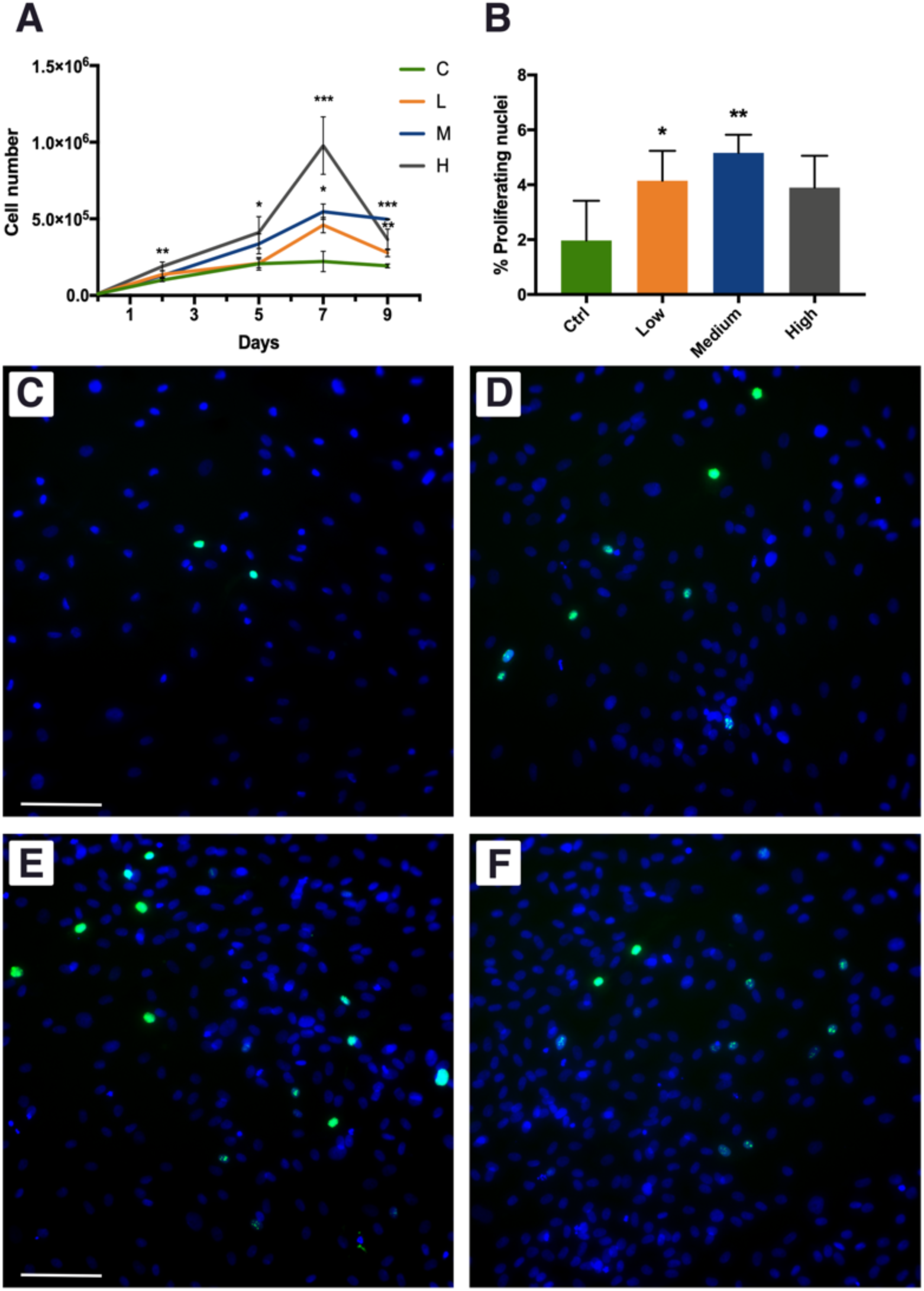
PrP affects hMSC proliferation. A) hMSC growth curves scored at different culture conditions: C (20% FBS), L (5×10^5^ platelets/ml), M (1×10^6^ platelets/ml) and H (1.5×10^6^ platelets/ml). B) Histogram representing Ki67 positive nuclei rate at the indicated media: Ctrl (20% FBS), Low (5×10^5^ platelets/ml), Medium (1×10^6^ platelets/ml) and High (1,5×10^6^ platelets/ml). Statistics: One-way Anova test. p > 0.05^*^, p > 0.01^**^, p > 0.001^***^ (n= 3). C-F) Immunofluorescence against Ki67 (green) on hMSCs at day 9 of different culture conditions: (C) Ctrl, (D) Low, (E) Medium, (F) High concentrations of PrP, nuclei were counterstained by DAPI (blue). Scale bars 150μm.

## Discussion

Nowadays, one of the most challenging tasks in skeletal muscle regenerative medicine is to find a suitable stem cells source for a large and easy cellular expansion, avoiding losing the myogenic potential (Errico et al., 2018). In fact, satellite cells, despite being the effective muscle stem cells, present a low isolation rate and can be kept in culture only for few passages without losing myogenic potentiality. This has prompted researchers to investigate other stem cell populations (Fuoco et al., 2016b, 2016a). However, despite the efforts of the last decades leading to the characterization of several stem cell populations with mesenchymal origin and promising myogenic potentiality, obtaining a large amount of human myogenic primary cells for tissue engineering or cell therapy approaches is still an unmet need (Y. et al., 2007; Skuk et al., 2014; Sacchetti et al., 2016). For this reason, in this work we evaluated how culture conditions can modulate different basic features of skeletal muscle-derived hMSCs, such as proliferation rate, myogenic potential efficacy and maintenance.

Here, we tested whether different growth media can directly influence hMSC - freshly isolated from human skeletal muscle biopsies - heterogeneity, expansion potentiality and differentiation capabilities. In particular, we employed different culture conditions comprising different media (Alpha-MEM and Cyto-grow) and different supplements (Human Serum and PrP). To monitor hMSC heterogeneity, we performed a time course analysis by cytofluorimetric assay, investigating mesenchymal stemness (CD90 positivity) and myogenic potential (CD56 positivity). The obtained results revealed that both media were able to maintain the cell population stemness for many doubling times (up to passage 9). Interestingly, while in standard medium (Alpha-MEM + 20% FBS) we observed the formation of two distinct subpopulations (single positive CD90^+^ and double positive CD90^+^/CD56^+^), in the rich medium a more homogeneous population double positive was formed and maintained, showing to directly promote the selection of the myogenic properties right from the early stages. Thus, in order to obtain a large number of myogenic stem/progenitor cells, the use of a rich medium - likewise Cyto-grow - would ameliorate remarkably the isolation, expansion and differentiation efficiency of hMSCs.

In parallel, we investigated the possibility to employ human derived serum as medium supplement in order to avoid animal derivatives and then guaranteeing completely autologous human cell condition, the so called “humanizing supplement” (Roberts et al., 2012; Shanbhag et al., 2017). Hence, we tested different human serum concentration (ranging from 5% to 20%), observing that at higher concentration (20%) the hMSC proliferation rate was about 5-fold greater than control (20% FBS). The medium dose (10%) had just a slight effect, while the lower concentration (5%) was comparable with the control in terms of proliferation rate. These results demonstrated the possibility to exploit human serum to ameliorate human cell proliferation rate and then to reduce the cellular expansion time. This strategy has a great clinical potential as, from the one hand, it avoids the use of animal-derived supplements and, form the other hand, lays the basis for the development of a culturing protocol that fulfill GMP regulations essential for human cell clinical application. Additionally, we have also investigated the impact of the human serum on hMSC myogenic capabilities, showing - upon 9 days of culture - a noteworthy enhancement of the differentiated myotubes in relation to the serum concentration, with a significant increment (up to 4-fold increase) of the fusion index at high concentration (20%).

Taken together these results suggest a strong direct correlation between human cell physiological processes, such as proliferation and differentiation, and medium supplementation with increasing doses of autologous serum.

The described outcomes referred to the human serum influence on hMSC behavior, led us to investigate whether the age of the serum donor could also affect cell growth, speculating that younger donor serum could have stronger effects than the older ones. In fact, it is well known and widely accepted than tissue regenerative potential, in relation to stem cell niche, milieu and availability, decreases with advancing age (Malecova, 2012; Tajbakhsh, 2013; Birbrair et al., 2014; Efimenko et al., 2014; Sousa-Victor et al., 2014; Rotini et al., 2018). Hence, to investigate the effect of serum donor age, we grouped the donors in three different groups in relation to their age: young (20-30 years old), adult (31-50 years old) and senior (51-65 years old). Unexpectedly, the proliferation rate did not show any significant influence by the different tested group sera, suggesting that donor age does not alter and/or modify human serum properties.

Finally, we tested PrP action on hMSCs to exploit another humanizing medium supplement largely used in clinical therapeutic approach for pathologies affecting musculoskeletal system (Roberts et al., 2012; Shanbhag et al., 2017). In this case, we observed that cell proliferation rate increases with the increase of PrP concentration, reaching the maximum at day 7 for the highest PrP concentration (approx. 4-fold increase compared to the control), further confirmed by Ki67 expression quantification (**Figure 4**). Nevertheless, at day 9 we noticed a dramatic decrease in the proliferation rate, with a more pronounced effect for the highest dose of PrP, probably due to the excessive over-confluence compromising cell health.

Hence, the presented results display a better effect of human serum on hMSCs compared with PrP, taking also into account the readiness of serum isolation and availability contrasting with the industrious PrP isolation and with the different formulations (Shanbhag et al., 2017).

In this study, we demonstrated the possibility to directly influence skeletal muscle-derived hMSC isolation and behavior improving cell proliferation rate and myogenic efficacy by formulating specific culture media. Moreover, we have shown that by exploiting human blood derivatives such as serum or PrP, one can achieve efficient and robust culture protocols for hMSC expansion and differentiation that do not require animal derived supplements and can be easily translated into the clinical scenarios fulfilling good manufacturing practice requirement.

## Acknowledgments

This study was supported by National Science Centre Poland (NCN) within the POLONEZ 3 Fellowship No. 2016/23/P/NZ1/03604 to M.C - this project has received funding from the European Union’s Horizon 2020 research and innovation programme under the Marie Skłodowska-Curie grant agreement No 665778. We thank FUNDACION ONCE for the cooperation and inclusion of people with disabilities supporting Dr. Carles Sanchez Riera research.

## Notes

### Competing Interest Statement

The authors have declared no competing interest.

